# A highly resolved integrated single-cell atlas of HPV-negative head and neck cancer

**DOI:** 10.1101/2025.03.02.640812

**Authors:** Lina Kroehling, Andrew Chen, Anthony Spinella, Eric Reed, Maria Kukuruzinka, Xaralabos Varelas, Stefano Monti

**Affiliations:** Bioinformatics Program, Faculty of Computing and Data Science, Boston University, Boston, Massachusetts, USA; Section of Computational Biomedicine, Boston University Chobanian & Avedisian School of Medicine, Boston, Massachusetts, USA; Department of Biochemistry and Cell Biology, Boston University Medical Center, Boston, MA, USA; Department of Medicine, Albert Einstein College of Medicine, Bronx, New York, USA; Department of Molecular and Cell Biology, Department of Translational Dental Medicine, Boston University Medical Center, Boston, Massachusetts USA; Department of Biostatistics, School of Public Health, Boston University, Boston, Massachusetts, USA

**Keywords:** Head and Neck Cancer, single-cell transcriptomics, single-cell atlas

## Abstract

Head and Neck Squamous Cell Carcinomas (HNSCC) are the seventh most prevalent form of cancer and are associated with human papilloma virus infection (HPV-positive) or with tobacco and alcohol use (HPV-negative). HPV-negative HNSCCs have a high recurrence rate, and individual patients’ responses to treatment vary greatly due to the high level of cellular heterogeneity of the tumor and its microenvironment. Here, we describe a HNSCC single cell atlas, which we created by integrating six publicly available datasets encompassing over 230,000 cells across 54 patients. We contextualized the relationships between existing signatures and cell populations, identified new subpopulations, and show the power of this large-scale resource to robustly identify associations between transcriptional signatures and clinical phenotypes that would not be possible to discover using fewer patients. We reveal a previously undefined myeloid population, sex-associated changes in cell type proportions, and novel interactions between CXCL8-positive fibroblasts and vascular endothelial cells. Beyond our findings, the atlas will serve as a public resource for the high-resolution characterization of tumor heterogeneity of HPV-negative HNSCC.

## Background

Head and Neck Squamous Cell Carcinomas (HNSCCs) arise from the mucosal epithelium lining of the oral cavity, larynx, and pharynx, with adult stem cells acting as cells of origin.[1,2] The two major subtypes are HPV-positive and HPV-negative HNSCC, the former more frequently occurring in the pharynx and associated with human papilloma virus infection, especially the HPV-16 strain [3,4], and the latter more frequently occurring in the oral cavity and larynx and associated with carcinogens in alcohol, tobacco and betel quid, among others.

Amongst all cancers, HNSCCs have the seventh highest incident rate, accounting for almost 7% of cancers worldwide, with 75% being HPV-negative.[3,5,6] According to the GLOBOCAN database, cancers of the lip and oral cavity are the leading cause of cancer death for men in India.[6] HPV-negative cancers tend to have higher mutational burdens than HPV-positive cancers and lower levels of immune infiltration in the tumor microenvironment (TME). Together, these factors result in clinicopathological differences and a significant difference in 5-year survival rates between the two subtypes, with a 75-80% rate for HPV-positive HNSCCs and a ∼55% rate for HPV-negative HNSCCs.[4,7–9] Consequently, the two subtypes have now been separated in the American Joint Committee of Cancer (AJCC) staging system and are commonly considered distinct diseases.[7,10]

Current treatments for the disease include radiation, chemotherapy, combination therapy, and immune checkpoint inhibitors (ICIs). However, response to treatment as well as mode of effective treatment varies greatly across individuals due to the high degree of heterogeneity both in the genetic profiles and plasticity of the cancer cells, as well as in the cell type composition of the TME, even within HPV-negative HNSCCs.[11,12] HNSCC patients have a poor overall response to ICI therapy, with less than 20% presenting a durable response.[11,13]

Significant progress has been made in characterizing the HNSCC TME. HPV-negative tumor epithelial cells are commonly characterized by tumor suppressor inactivating mutations such as TP53, CDKN2a, and PTEN, and amplifications in CCND1, which together enable the progression of the disease.[4,14] Previously identified transcriptional signatures in epithelial cells include response to hypoxia, stress, epithelial-to-mesenchymal transition (EMT), and partial EMT (pEMT), a molecular phenotype in which tumor cells de-differentiate to become more mesenchymal-like and acquire enhanced migratory capabilities.[15] Additionally, immune and stromal cells present in the TME play a major role in affecting tumor growth through TME interactions. For example, cancer-associated fibroblasts (CAFs) have been shown to affect tumor cell migration by remodelling the extracellular matrix.[15] Natural killer (NK) and CD8 T cells have been found to take on effector phenotypes and kill tumor cells, whereas myeloid-derived suppressor cells (MDSCs) and Tregs secrete cytokines that suppress the immune system.[4] Further delineation of these complex dynamics in the non-HPV TME is critical for discovering new potential avenues of therapeutic intervention that may drive heterogeneous patient responses to ICI.

Single-cell RNA-sequencing (scRNA-seq) has emerged as the technology of choice to profile the individual transcriptomes of tumor cells and the surrounding TME. scRNA-seq enables the characterization of tumor heterogeneity within and across patients through the identification of *cell types* – here defined as categories of cell that perform a specific function (e.g., T cell, Fibroblast), and specific *cell states* – here defined as transient molecular changes in gene expression in response to endogenous and exogenous cues that multiple cell types can share (e.g., dysfunction, proliferation).[15–18] While single-cell HNSCC datasets have produced invaluable findings, given the relatively small sample size of the individual studies, and the relatively small number of publicly available single-cell HNSCC datasets, we hypothesized that their integration and combined analysis would facilitate contextualization of findings from individual studies, and provide increased statistical power to identify novel associations between molecular and clinical phenotypes.[19] HNSCC atlas’ have been developed, however the focus has been on a single cell type[20,21], contrasting the HPV-positive and HPV-negative subtypes16, or on the stepwise progression of the disease[22,23], leaving a need for the comprehensive characterization of the HPV-negative subtype.

To this end, we have integrated six publicly available datasets [Puram et al.[15], Cillo et al.[24], Kurten et al.[25], Peng et al.[26], Choi et al.[23], Quah et al.[27]] for the high-resolution characterization of the cells in the TME of HPV-negative HNSCC. Creation of the atlas allowed us to perform four types of analyses. **Contextualization**, in which we taxonomically organize different cell types and identify how closely related previously identified populations are to each other, and perform cluster co-occurrence analysis to identify cell populations whose changes in proportions are associated across patients. **Reconciliation**, in which previously described populations with the same characteristics but differently labelled in the individual studies are assigned a consistent nomenclature. **Identification and characterization of new subpopulations and interactions,** in which we leverage the increased number of cell profiles in the atlas to identify and annotate novel cell types. **Association with clinical phenotypes**, in which the increased sample size of the atlas increases the statistical power to detect significant associations between cell type composition changes, cell type and state signatures, and clinical phenotypes such as stage and sex.

In summary, we leverage published data to create and annotate an HPV-negative HNSCC transcriptomic atlas comprised of over 50 patients and 230,000 cells that may serve as a public resource. The atlas integrates six existing datasets to produce a comprehensive landscape of HNSCC tumors and their microenvironment, supporting novel findings, providing a reference for generating and contextualizing new hypotheses, and connecting molecular phenotypes to clinical end points.

## Results

### Generation of an integrated atlas

Six publicly available datasets containing human HPV-negative HNSCC tumors were combined, allowing for the profiling of 54 patients and over 232,015 cells (Table 1). Aggregated patient metadata is supplied in Supplementary Table 1. The integrated UMAP shows both mixing of datasets and maintenance of cell classification purity, with cells assigned to a specific cell type pre-integration clustering together post-integration (Fig 1a,b). Clusters contained cells from different sexes, sites, ages, stages, and datasets (Supplementary Fig 1a, b). Despite differences in biology between different anatomical sites28, cells from the larynx and pharynx are interspersed with the majority of cells which come from the oral cavity (Supplementary Figure 1a). Post-integration, some patient-specific epithelial clusters remained, and likely represent tumor-specific clones that expanded within an individual (Supplementary Fig 1c). Canonical marker genes for broad-level cell types showed specificity in their expression patterns (Fig 1c). The datasets varied greatly in both the number of cells recovered from each patient, and the cell types recovered from each patient, emphasizing the heterogeneity that exists in the tumor microenvironment between patients (Fig 1d).

**Figure 1:**
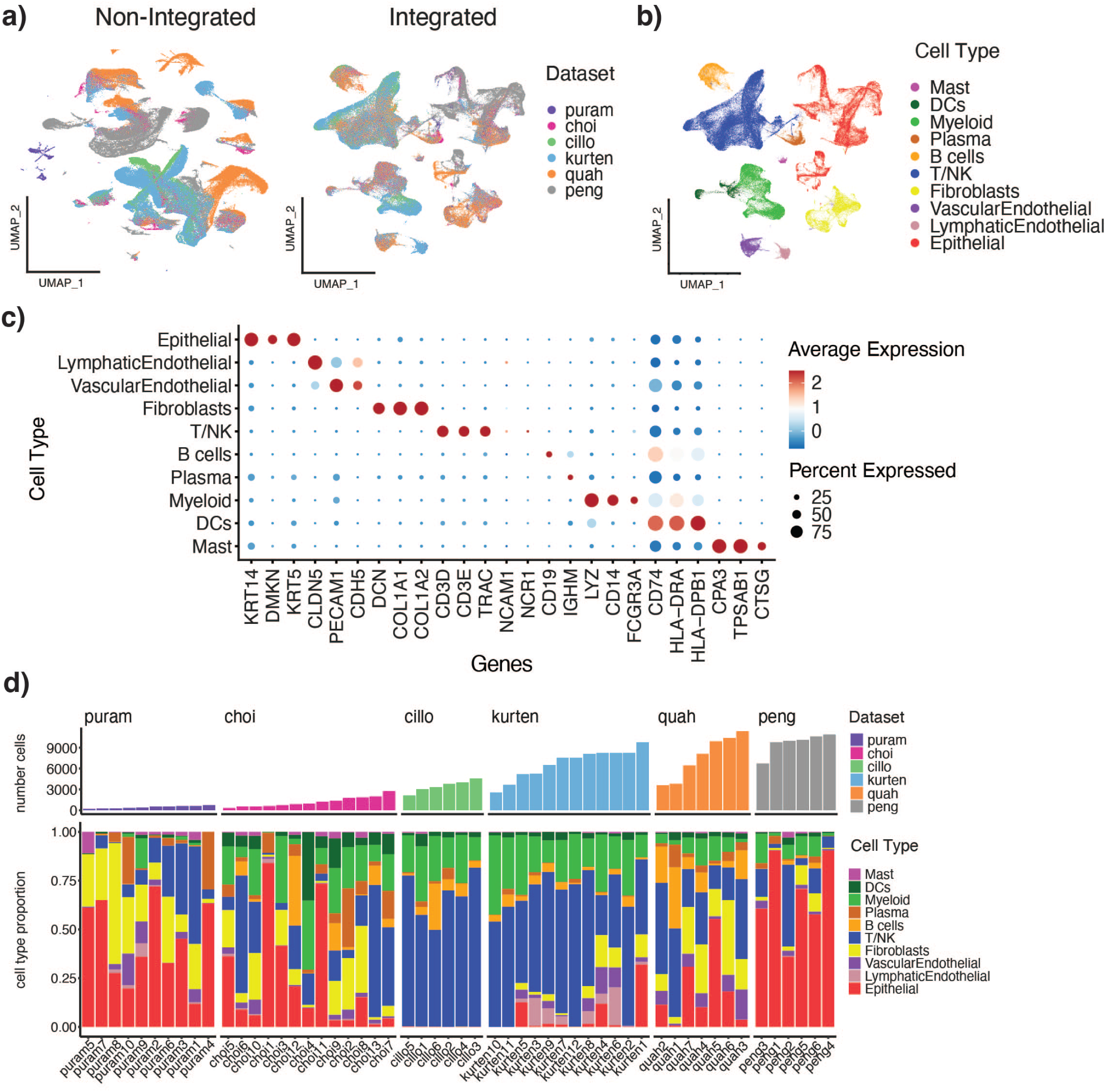
Creation of integrated HPV-negative scRNAseq atlas reveals inter– and intra-tumor heterogeneity. a) Merged but nonintegrated UMAP (left) and integrated UMAP (right) colored by dataset. b) Integrated UMAP colored by cell type. c) DotPlot showing average expression of canonical marker genes by cell type. d) Distribution of cell types per patient (bottom), and total number of cells per patient (top).

**Table 1:** Summary of datasets used in atlas.

| Dataset | GEO | # Patients (HPV-) | # Cells | Sorting | Age Range | Sex | LNM (primary/LNM) | T Stage |
| --- | --- | --- | --- | --- | --- | --- | --- | --- |
| Puram | GSE103322 | 10 | 4199 |  | 41-88 | 4/6 | 5/5 | 1/3/2/4 |
| Kurten | GSE164690 | 12 | 80523 |  | 35-80 | 5/7 | 12/0 | 1/2/7/2 |
| Cillo | GSE139324 | 6 | 20692 | CD45+ only | 43-66 | 1/5 | 6/0 | 1/1/1/3 |
| Peng | GSE172577 | 6 | 57984 |  | 32-72 | 2/4 | 6/0 | 0/2/2/2 |
| Quah | GSE225331 | 7 | 53459 |  | 18-70 | 2/5 | 0/7 | 0/0/4/3 |
| Choi | GSE181919 | 13 | 15158 |  | 53-77 | 4/9 | 13/0 | 2/8/0/3 |
| **Total** |  | **54** | **232015** |  | **18-88** | **18/36** | **42/12** | **5/15/16/18** |

### Separation of Immune and Non-immune Compartments

Because some datasets sorted cells before integration, (Cillo et al. contained only CD45+ cells), we split the atlas into the immune and non-immune compartments for downstream analyses. To this end, we first performed global clustering on the entire integrated dataset at resolution 0.4 and identified major cell compartments within. The non-immune compartment was defined as the union of cells classified as Adipocytes, Endothelial cells, Epithelial cells, Fibroblasts, and Muscle cells. The immune compartment was defined as the union of cells classified as T cells, NK cells, B cells, Immune (a broad label containing myeloid cells), and Mast cells. The final immune and nonimmune subsets were reclustered at a resolution of 0.8 and 0.4, respectively (Supplementary Fig 2a).

### Immune Compartment

Within the immune compartment, we identified 30 clusters, which we classified into five broad cell types: myeloid, T/NK cells, B cells, Plasma cells, and Mast cells, based on a consensus annotation that integrated singleR classification and the use of canonical marker genes (Fig 2a,b). Expression of markers CD14 and CD16 (FCGR3A) separated monocytes and macrophages within the myeloid group, while markers CD4, CD8A, and CD160 clearly separated CD4, CD8 and NK cells within the T/NK cell cluster (Fig 2b). We also identified plasma cells (IGHG1-positive), B cells (CD20/MS4A1-positive), and Mast cells (TSAB1-positive). Enrichment of marker genes and signatures revealed clusters of cDCs (clusters 14, 25, 26), pDCs, (19), macrophages (5, 7), and monocytes (2, 17, 24) (Supplementary Figure 3a,b).

**Figure 2.**
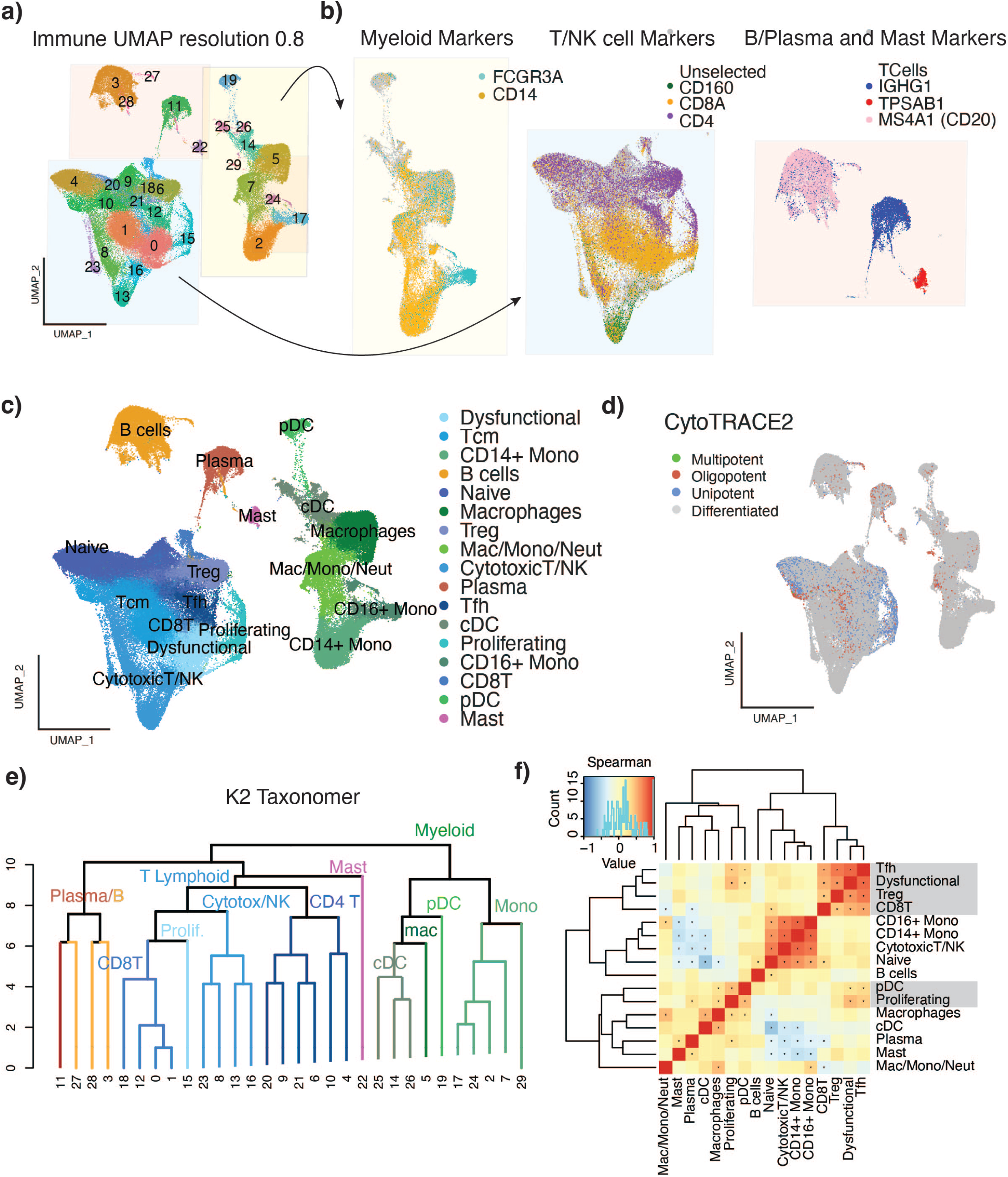
Characterization of cell types in the immune microenvironment. a) Immune compartment UMAP clustered at resolution 0.8. b) Identification of broad cell types by expression of marker genes. FCGR3A and CD14 expression on myeloid cells (left), CD160, CD8A and CD4 expression on T and NK cells (center), and IGHG1, TPSAB1 and MS4A1 expression on B cells, plasma cells, and mast cells (right). Cells highlighted are those expressing more than the average gene expression across the immune compartment for the given gene. c) UMAP of immune compartment with cluster names based on signature enrichment. d) UMAP of immune compartment colored by CytoTRACE2 potency classifications. e) K2 Taxonomer dendrogram of immune compartment clusters from resolution 0.8. f) Immune cluster-cooccurrence. Correlation calculated using spearman correlation * P < 0.05 and R >= 0.01.

#### T cell annotation reconciles previous findings, identifies rare sub-types and novel associations of cellular programs with clinical phenotypes

To resolve cell subtypes at a higher resolution within the broad T/NK cell group, we scored T cells using curated signatures and evaluated their enrichment at the cluster level (Supplementary Figure 3c). We identified clusters of naïve T cells, T follicular helper cells (Tfh) regulatory T cells (Tregs), and central memory T cells (Tcm) within the CD4 compartment, and cytotoxic, central memory, and dysfunctional cells within the CD8 compartment, as well as populations of NK and proliferating T cells. (Figure 2a-c, Supplementary Figure 3d)

##### Identification of CD4 T-cell subtypes: Naïve, Tregs, and T follicular helper cells (Tfh)

Naïve T cells were significantly enriched in clusters 4, 9, 10, 20 and 21, while Tregs were more abundant in cluster 6, and Tfh cells in cluster 12, all of which expressed CD4 (Supplementary Figure 3c). Across the published studies the naïve population has been referred to by a variety of names: naïve (Choi[23]), CD4 (Kurten[25]), naïve-like (Quah[27]), conventional T-helper (Puram[15], Cillo[24]), and TCF7+ cells (Peng[26]), despite expressing a consistent set of genes (IL7R, LTB, CCR7, SELL, TCF7).

##### Natural Killer (NK) and CD8 T cells fall on a trajectory from cytotoxicity to dysfunction

Next, we focused on CD8 and NK cells. Clusters 13 and 16 displayed up-regulation of CD160, previously shown to be expressed in peripheral blood NK cells[29], and enrichment of the NK cell signature (Supplementary Figure 3c). Clusters 0, 1, 8, and 23 were identified as CD8+ T cell clusters. Each cell was further annotated by *dysfunctional* and *cytotoxic* scores[30], and an overall effector capacity score was obtained by subtracting the dysfunctional score from the cytotoxic score, with a higher score indicating higher levels of cytotoxicity and less dysfunction. The NK cells (clusters 13, 16) showed the highest cytotoxic score, reflecting their known cytotoxic ability, followed by CD8+ T cell clusters 0, 8, and 23, with CD8+ T cell cluster 0 also displaying the highest dysfunctional score, reflecting a multi-state phenotype characterized by expression of both cytotoxicity and dysfunction programs.

##### The balance of T cell cytotoxicity and dysfunction is associated with tumor stage

We investigated the T cell effector capacity score’s association with stage. Using a linear mixed effect model to regress the effector capacity score (cytotoxic – dysfunctional) on stage, we found the score to be significantly positively associated with stage 1 (p-value=0.0201), and significantly negatively associated with stage 2 (p-value=0.0408), suggesting a cytotoxic T cell response at the earliest stage, becoming exhausted at later stages (Supplementary Figure 3e). The same analysis performed within the individual datasets showed the estimates to vary greatly in effect size and direction and mostly failing to achieve significance, except for in the Kurten dataset, highlighting the utility of the large *n* in the atlas to identify associations with clinical features.

#### Contextualization of Immune Cell Populations

We next sought to elucidate the relationship between cell-type annotated clusters in the immune compartment (Fig 2c).

##### Immune cell potency

CytoTRACE2 is an interpretable AI method for predicting cellular potency from scRNA-seq data. We applied this tool to the analysis of the immune compartment and found that most immune cells were in a fully differentiated state (Fig 2d). The naïve T cells and proliferating T cells were classified as unipotent or oligopotent, as they can differentiate into T cell subsets such as cytotoxic T cells or Tregs. Interestingly, some T cells were also annotated as unipotent. These results are consistent with previous studies showing that Tregs are highly plastic and may switch between states in response to different stimuli.[31,32]

##### Contextualization of immune cell populations

Application of K2Taxonomer (K2T) to the immune cell compartment identified the taxonomic relationships between the clusters therein[33] (Fig 2e). The top-level partition separated the myeloid and lymphoid subsets, except for the Mast cells which clustered within the lymphoid clade, likely due to negative expression of some myeloid markers such as LYZ and LST1. (Supplementary Fig 4a). The four monocyte clusters were grouped within a clade (2, 7, 17, 24, and 29). Cluster 5 (macrophages), was most closely related to cDC clusters 14, 22, and 26, due to elevated expression of complement genes in both the macrophages and cDCs compared to the rest of the immune cells (Supplementary Fig 4b). The T-lymphoid and NK cells were grouped, and separated into clades that can be divided by cell states: naïve (4, 9, 10, 20, 21), cytotoxic (8, 13, 16, 23), and dysfunctional (0).

Concurrently, cluster co-occurrence analysis showed that naïve, cytotoxic, NK, and monocytes were highly correlated in their proportional changes across patients, and were generally associated with better prognosis. Tregs, Tfh, and dysfunctional cells were highly correlated and associated with worse survival (Fig 4e), while also correlated with pDCs. Naïve cells were significantly positively correlated with B cells, a result consistent with Peng et al., where TCF7+ T cells (naïve) were shown to be distributed in and around tertiary lymphoid structures (TLSs) made up of CD20+ B cells.[26]

#### Macrophage annotation identifies known and novel subtypes

As recent studies have identified new polarization states of macrophages, we leveraged the atlas’ large sample size to, first, recapitulate these states, and then further characterize the macrophages at high resolution. Clusters 5 and 7 were identified as potential macrophages due to the expression of C1QB and macrophage cell type signatures and a lack of enrichment for classical monocyte signatures (Supplementary Fig 4b). These cells were reclustered at resolution 0.3 to identify six subpopulations therein (Fig 3a).

**Figure 3:**
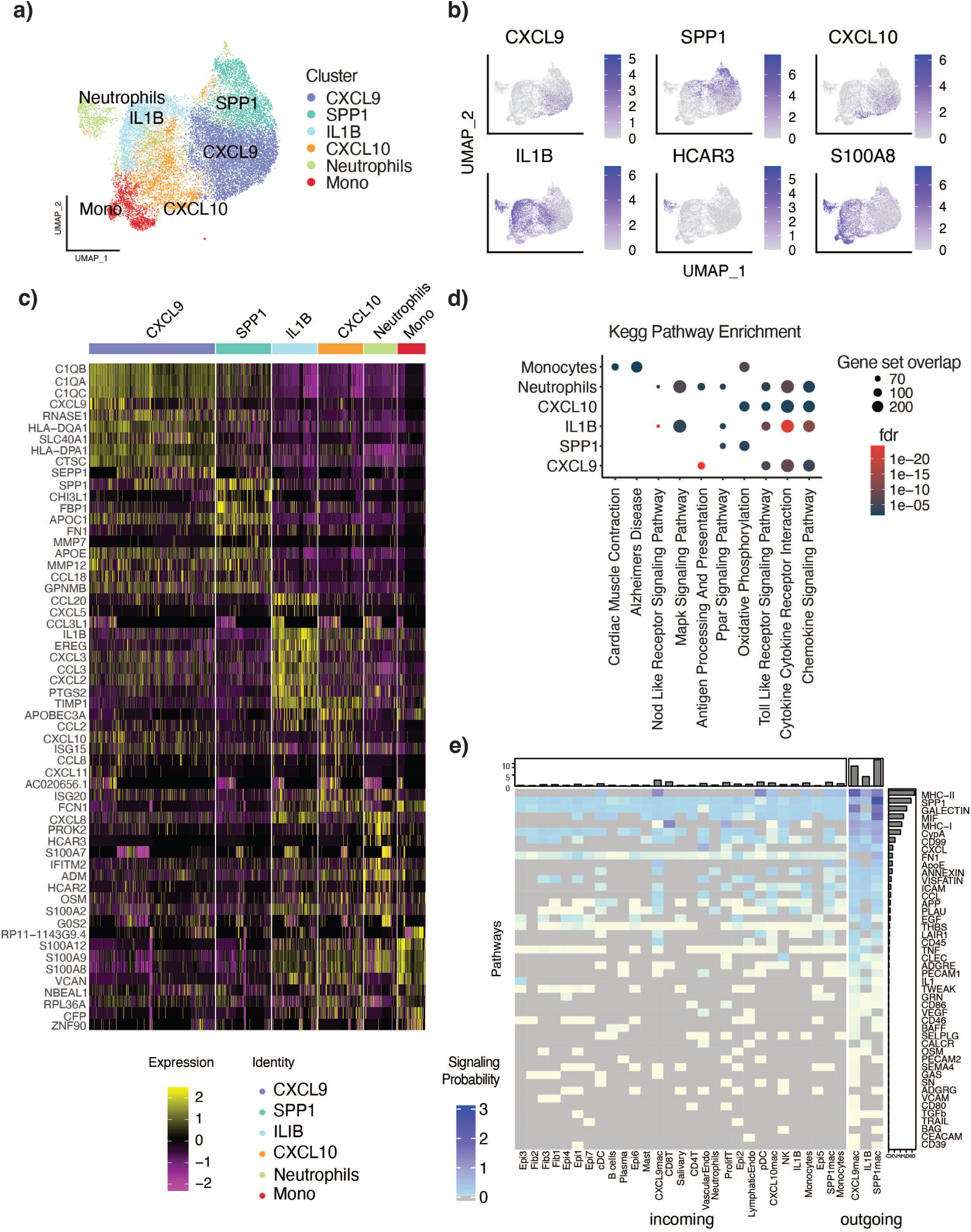
Identification of macrophage subclusters and signaling pathways. a) UMAP of immune compartment clusters 5 and 7 reclustered at resolution 0.3. Six clusters were identified. b) UMAPs of marker gene expression for the six clusters, CXCL9, SPP1, CXCL10, IL1B, HCAR3, and S100A8 from left to right. c) Heatmap of top 10 marker genes per cluster. d) Kegg signature enrichment for macrophage subclusters based on signature genes using hypergeometric test. e) Cell cell communication analysis by CellChat showing pooled signaling probability across pathways. Signals shown are sent from the CXCL9, SPP1, and IL1B, clusters (outgoing) to the other clusters (incoming).

**Figure 4:**
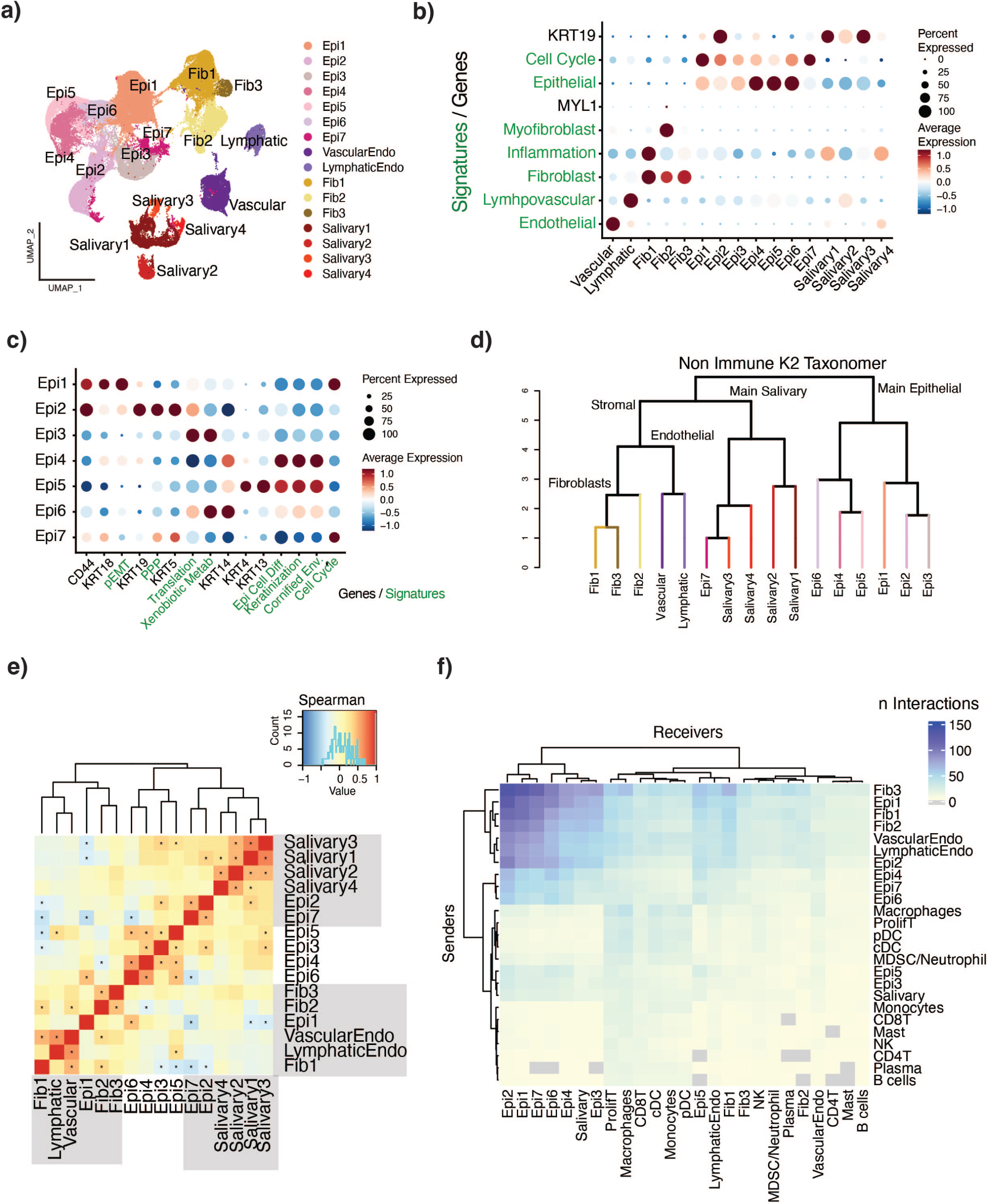
Non-Immune compartment characterization and contextualization. a) UMAP clustered at resolution 0.4 for a total of 16 clusters. b) DotPlot showing expression of marker genes (black) and cell type signatures (green) across all non-immune cells. c) DotPlot showing expression of marker genes (black) and signatures (green) across seven epithelial clusters. d) K2Taxonomer organization of clusters. e) Cluster co-occurrence analysis by spearman correlation. * P < 0.05 and R >= 0.01. f) Cell-cell communication analysis by CellChat. The heatmap is showing the pooled number of predicted interactions per patient in a pairwise manner between each cluster.

Interrogation of the top genes enriched for each cluster revealed two clusters, SPP1-positive and CXCL9-positive macrophages, showing high expression of macrophage marker genes C1QB, CD4, and CD163 (Fig 3b,c). The CXCL9 cluster was further uniquely defined by high expression of CXCL9 and enriched for antigen presenting-related genes (Fig 3b,d). The SPP1 cluster was defined by high expression of SPP1 and enriched for metalloproteinases (MMPs), coincident with a relative dearth of Kegg or Hallmark terms (Fig 3c,d). These macrophage clusters were distinguished by CXCL9 and SPP1 rather than by the canonical M1 and M2 markers (Supplementary Fig 4c), lending further support to an alternative macrophage categorization by polarization states in HNSCCs recently described by Bill et al.[34]

The remaining four clusters were negative for macrophage marker genes C1QB, CD4, APOE and CD163 (Supplementary Fig 4a). We identified a CXCL10-positive cluster that also over-expressed IFIT genes and CCL2 and was enriched for interferon alpha and gamma responses (Fig 3c,d). This cluster is similar to a population described by Zhang et al. in inflammatory conditions, expressing genes such as IDO1 and GBP1, and stimulated by IFN-γ and TNF-α.[35] Additionally, we identified a cluster not previously described in HNSCC with characteristics of myeloid-derived suppressor cells (MDSCs), particularly characterized by high expression of inflammatory cytokines including IL1β, but negative for the macrophage markers SPP1 and CXCL9. Finally, a cluster of Neutrophils was also identified.

The IL1β cluster showed similarity with VCAN– and EREG-expressing tumor-associated macrophages (TAMs) associated with angiogenesis identified in a pan-cancer macrophage atlas.[36] This cluster was also found to be enriched for inflammatory genes, including the aforementioned IL1β, which has been shown to promote drug resistance through induction of ICAM1 on tumor cells[37], IL6 and IL10, which have been shown to promote immunosuppression through induction of Tregs[38], cytokine-cytokine receptor interactions, and EMT (Fig 3c,d). This population shares some transcriptional similarity with HNSC TAM populations previously described as M2 macrophages[39]. However, we observed key differences in the expression of several key M2 markers such as a lack of CD163 expression, the absence of complement gene expression, and the lack of ARG1 expression.[40] Comparison of the IL1β cluster’s cell-cell interactions with those of the SPP1 and CXCL9 clusters identified Thrombospondin (THBS) signaling as specific to this cluster, as well as increased tumor necrosis factor (TNF) and IL1β signaling (Fig 3e). IL1β and TNF-α secretion by SPP1-positive macrophages has been described and attributed to progressing HNSCC[41], but the atlas highlights the presence of this IL1β-positive SPP1-negative population as a discrete population that may be involved in unique signaling pathways contributing to tumor progression and drug resistance.

The Neutrophil cluster appeared to be neutrophil-like due to its lack of HLA-DRA, HLA-DRB1, and CD74 expression, and its expression of IFITM2, S100A8, PROK2, HCAR3, CMTM2, and CSF3R, among others.[42,43] A gene set enrichment analysis showed Neutrophils to be enriched for cytokine-cytokine receptor interactions (Fig 3d). Cell type classification by HPCAD and Blueprint databases supported these conclusions (Supplementary Fig 4d).

#### Non-immune compartment

Clustering analysis of the non-immune compartment based on Seurat at resolution 0.2 identified 16 clusters, including separate clusters of vascular and lymphatic endothelial cells, three fibroblast clusters, seven tumor epithelial cell clusters, and four salivary cell clusters, as expressing canonical marker genes and cell type signatures (Fig 5a-c4a,b).

**Figure 5.**
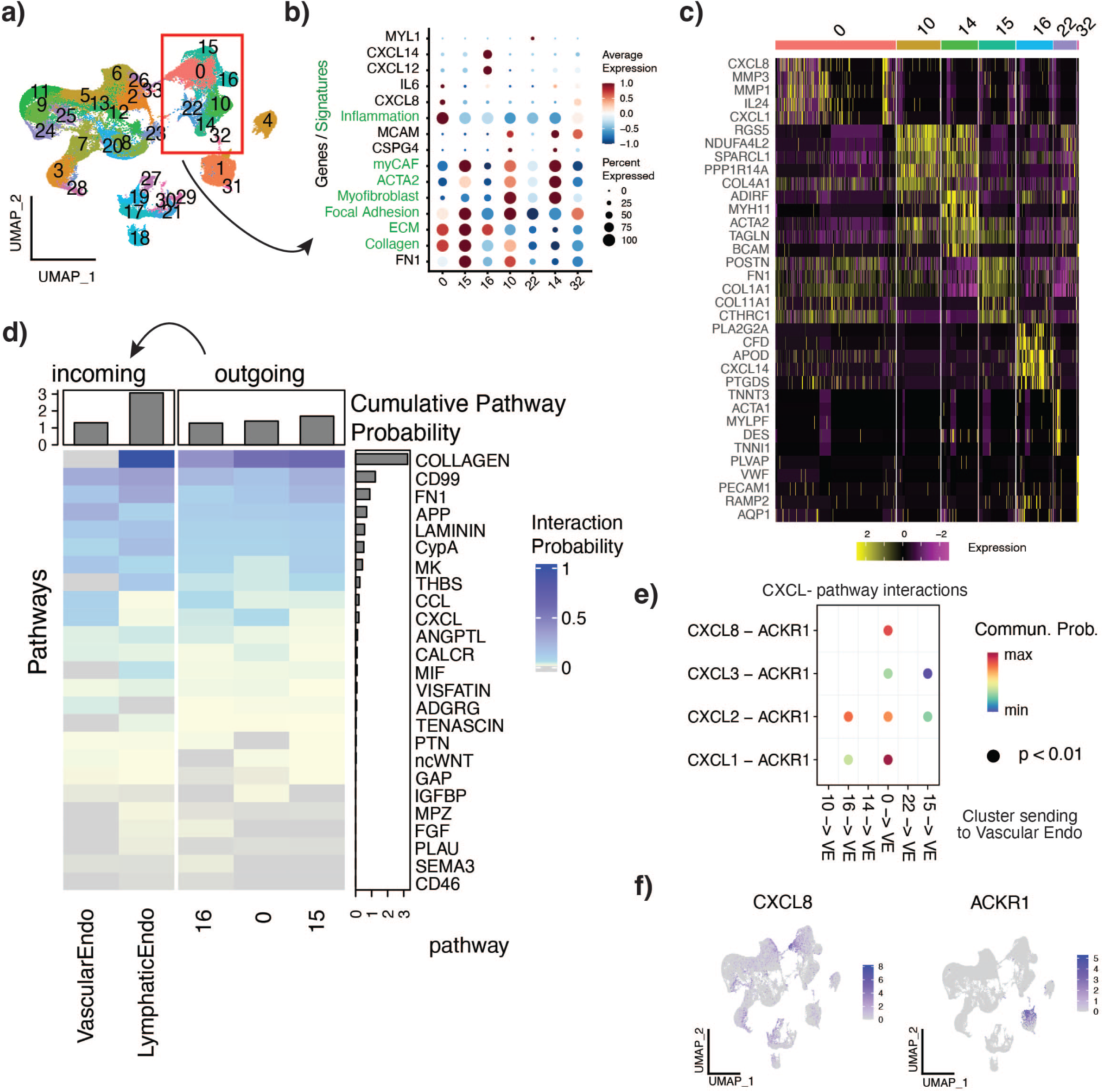
Fibroblast classification and signaling. a) UMAP clustered at resolution 0.7 for a total of 34 clusters. b) DotPlot showing expression of marker genes (black) and cell type signatures (green) across fibroblast clusters. c) Heatmap of top five DE genes per fibroblast cluster. d) Cell-cell communication analysis by CellChat showing pathways involved in sending signals from fibroblast clusters 0, 15, and 16 (outgoing) to the vascular and lymphatic endothelial cells (incoming). e) Specific ligand and receptor pairs involved in CXCL pathway interactions occurring between fibroblast clusters 0, 15, and 16 and the vascular endothelial cells. f) Expression of CXCL8 and ACKR1 in nonimmune compartment.

##### Epithelial cells segregate on a differentiation continuum

Annotation of the epithelial compartment highlighted a differentiation trajectory from basal cells to squamous cells, with cluster Epi1 being the least differentiated, progressing along clusters Epi6, Epi5, and Epi4 (most differentiated) (Fig 4c). Epi1 was enriched for CD44, a marker of CSCs[44], and KRT18, expressed in progenitor cells and malignant epithelial cells[45]. The pEMT program was also enriched, further suggesting that this cluster was likely to be less differentiated with plastic characteristics. Epi4, Epi5, and Epi6 were found to be enriched for keratinization, cornification, and epithelial cell differentiation (Fig 4c). Epi6 showed high expression of KRT5 and KRT14, characteristically co-expressed in basal cells[46], indicating that this cluster was at an early stage of the differentiation trajectory. Epi4 showed high expression of KRT13, KRT4, and SPRR3, genes characteristic of suprabasal cells.[47]

To further investigate the differentiation status of these cells, we applied CytoTRACE2 to the non-immune compartment to predict each cell’s potential to differentiate. The cytoTRACE2 potency scores are a continuous measure of developmental potential, and range from 0 (differentiated) to 1 (totipotent), with thresholds that distinguish the differentiated, unipotent, oligopotent, multipotent, pluripotent and totipotent categories.[48] Most of the epithelial cells were predicted to be in an oligopotent state whereas the rest of the cells in the tumor microenvironment were predicted to be fully differentiated (Supplementary Fig 5a). Within the epithelial clusters, Epi1 cells had the highest CytoTRACE2 potency scores, followed by Epi6, whereas Epi4 and Epi5 had lower scores, suggesting that potency was inversely correlated with differentiation. Interestingly, some of the cancer-associated fibroblasts (CAFs) were also predicted to be oligopotent, (defined as a lineage-restricted potential to differentiate into 2-3 downstream cell types), which may reflect their ability to be reprogrammed.[49]

##### Some epithelial clusters are patient-specific

Hierarchical organization of the non-immune compartment by K2Taxonomer first partitioned the cells into (mainly) stromal and epithelial groups, with epithelial cluster pairs Epi4 and Epi5 grouped together, further supporting the previously identified differentiation trajectory (Fig 4d). Interestingly, Epi7 was grouped within the stromal partition. When assessing the distribution of cells from each patient across each epithelial cluster (Supplementary Fig 5b), we found that clusters Epi2, Epi3, and Epi5 consisted primarily of cells from individual patients, and we observed specific enrichment for pathways not associated with other epithelial cells, such as the pentose phosphate pathway and xenobiotic drug metabolism (Epi2), and translation (Epi3), with Epi7’s signature consistent with a cycling population of cells (Fig 5c). Interestingly, Epi2 also expressed Krt19, which has been shown to reprogram CSCs in breast cancer to a more-drug-sensitive state.[50] Due to the specific enrichment patterns and patient composition of these clusters, they likely represent specific tumor-clone evolving populations.

##### Copy Number Variation identifies tumor subclones

To estimate the copy number variation status of the epithelial cells, we computed an aggregated malignancy score per cell from CNVs identified via inferCNV, (Supplementary Fig 5c), defined as the total amount of CNVs in a cell, regardless of direction (amplification or loss), with a higher number corresponding to a higher number of alterations, and a lower number corresponding to a cell with less CNVs. Different clusters varied significantly in their malignancy score distributions, with patient-specific clusters Epi2 and Epi3 having the highest and lowest malignancy scores within the epithelial compartment, respectively.

##### Less differentiated cells are more likely to exist on the tumor edge and interact with the TME

Cluster co-occurrence analysis (Fig 4e) showed that cluster Epi 1 was more strongly associated with the stromal clusters than with the other epithelial clusters, although no significant correlation was seen between Epi 1 and the stromal cells. To further investigate the hypothesis that Epi1 cells have higher interactions with the stromal cells and may be differentially spatially located within the tumor, we leveraged an external dataset, Arora et al[51], which spatially profiled HNSC tumors and derived signatures of a tumor leading-edge and of a tumor core. Projection of the cell profiles onto these signatures showed that Epi1 cells were enriched for the leading-edge signature, whereas Epi4, and to a lesser degree Epi5, were enriched for the tumor core (Supplementary Figure 5d). This is consistent with previous findings, which showed epithelial cells expressing pEMT markers to be located at the tumor front, facilitating their interaction with CAFs.[15] Additionally, Epi1 is strongly expressing TGFBI, which is secreted and becomes integrated into the ECM, and is associated with aggressive cancers.[52,53] A cell cell communication analysis of the total number of pairwise interactions between clusters showed a high number of interactions originating from all fibroblast and endothelial clusters, as well as from Epi1 and Epi2, towards many of the Epithelial clusters (Fig 4f).

##### Detailed characterization of epithelial cells

In order to further characterize the tumor cells, we subset and reclustered the epithelial cells at a resolution of 0.4 and identified 18 distinct clusters (Supplementary Fig 5e). Marker genes were shared across multiple high-resolution clusters, such as ACKR1C3 for clusters 0 and 4, suggesting that the initial lower resolution clustering was sufficient to capture the most salient cell differences (Supplementary Fig 5f). However, some fine distinctions between high-resolution clusters could be identified, such as the upregulation of a specific set of genes in cluster 14 and 15 (VAPA, CAPZA2, VIM) that were expressed at lower levels in clusters 0, 4, suggesting that clusters 14 and 15 are the most malignant subclusters of the low resolution cluster Epi1. Gene set enrichment analysis using the hallmarks compendium and signatures from Puram et al.[16] revealed pathways co-enriched in multiple epithelial cell clusters (Supplementary Fig 5g). These included cholesterol homeostasis, known to be important for proper keratinization, and epithelial cell differentiation, both enriched in clusters 7, 8, 9 and 13.[54] Enrichment of translation, Notch signaling, apoptosis and the p53 pathway was observed in clusters 3, 11, and 17, suggesting Notch signaling is driving changes in these clusters, as notch signaling has been shown to interact with the p53 pathway to inhibit apoptosis and drive aberrant translation.[55,56] Enrichment of hypoxia, Tgfβ and Il6 signaling was observed in clusters 10, 12, 15, and 16, with hypoxia previously shown to induce Tgfβ and IL6 signaling in cancer.[57,58]

##### Characterization of atlas fibroblast clusters

Next, we focused on CAFs, a heterogeneous population of cells that represents an important component of the tumor microenvironment and plays several key roles in cancer progression.[15,59–61]

Clustering of the stromal and epithelial compartment at a resolution of 0.7 identified seven populations of fibroblasts (Figure 5a), which we scored using known fibroblast genes and signatures. Signatures were defined for each cluster using a log2 fold change cutoff > 1 and FDR < 0.05 (Supplementary Table 4).

Interrogation of marker genes and signatures for each cluster showed clusters 0 and 15 to represent extracellular matrix related CAFs due to their enrichment for collagen and ECM signatures (Fig 5b,c). Cluster 16 was also enriched for the ECM signature but also expressed immune-related genes such as CXCL12 and CXCL14. Clusters 10, 14, 22, and 32 were identified as myofibroblast-like pericytes, as they showed high expression of smooth muscle actin (ACTA2, a myofibroblast marker) and pericyte marker genes (Fig 5b,c). Cluster 0 expressed a separate set of immune related genes, including CXCL8, CXCL1, MMPs and IL24, and was enriched for inflammation (Fig 6b-c). Cluster 22 likely consisted of some myocyte muscle cells due to their expression of myosin (MYL1).

**Figure 6.**
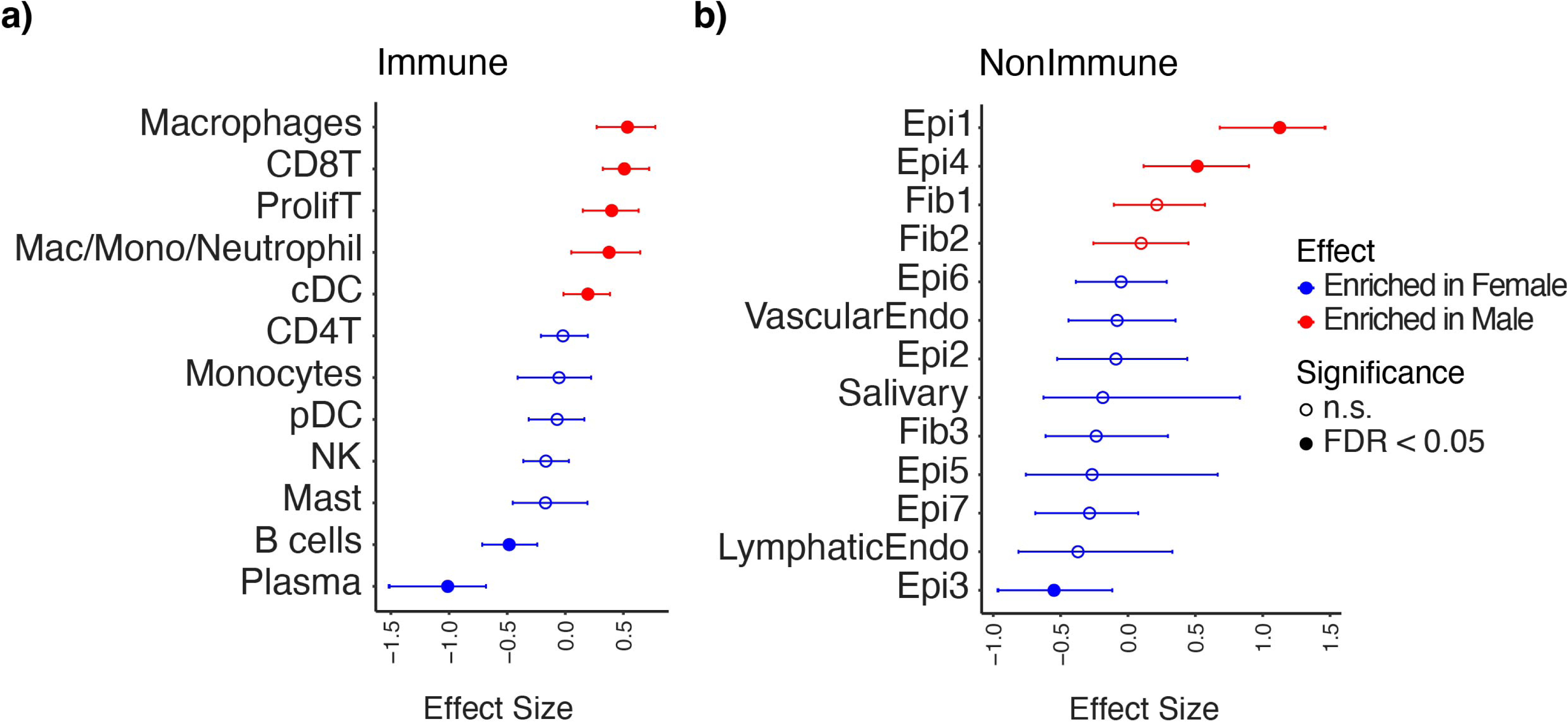
Sex-specific cell type proportion changes. a) Cell type proportion changes identified by sccomp within the immune compartment. b) Cell type proportion changes identified by sccomp within the nonimmune compartment.

To elucidate how the atlas fibroblast clusters overlapped with the CAF populations identified in the originating studies, we scored the atlas fibroblast clusters using CAF gene signatures reported by each study (Supplementary Fig 6a), toward a reconciliation of the nomenclature adopted by different studies to define the same populations (Fig 6a, Table 2).

**Table 2:** Fibroblast nomenclature reconciliation.

|  | Cluster |  |  |  |  |  |  |
| --- | --- | --- | --- | --- | --- | --- | --- |
| Dataset | 0 | 15 | 16 | 14 | 22 | 10 | 32 |
| Atlas | CXCL8-CAFs | ECM-CAFs | Imm-CAF (CXCL8) | Myofibroblasts-1 | Myofibroblasts-2 | Myofibroblast-ECM | Myofibroblasts-3 |
| Puram | CAF1 | CAF1 | CAF2 | Myofibroblasts | NA | CAF1 Myofibroblasts |  |
| Choi | cf4 | cf4 | cf2,cf4 | cf3 | Myocytes | cf3 | cf3,cf4 |
| Kurten | CAF | CAF | Elastic | Pericytes | Pericytes | Pericytes | Pericytes |
| Quah | CAF | CAF | CAF | Myofibroblast | Myofibroblast | Myofibroblast | CAF,Myofibroblast |
| Peng | No Fib analysis |  |  |  |  |  |  |
| Cillo | No CD45- cells |  |  |  |  |  |  |

Cells within Cluster 15, the ECM-related CAFs, were generally termed CAFs by most studies and were the predominant CAF type across studies.[15,23,25,27] Cluster 0 and 16, the two immune-related populations varied in their terminology, including ‘elastic’ and ‘resting’.[15,25] Clusters 10, 14, 22 and 32 were found to be myofibroblasts, and were termed either myofibroblasts or pericytes by most studies.[15,23,25,27]

Due to the high expression of numerous CXCL genes among the CAFs in clusters 0, 15, and 6, and the observed high co-occurrence of fibroblasts and endothelial cell clusters (Fig 4e), we examined the role of fibroblast signaling to the endothelial cells. Notably, fibroblasts were found to send collagen signals to the lymphatic endothelial cells, but not to the vascular endothelial cells. (Fig 5d) Cluster 0 showed increased signaling strength to the vascular endothelial cells via the CXCL pathway (Fig 5d), through both increased strength in CXCL1-ACKR1 interaction and a CXCL8-ACKR1 interaction (Fig 5e). Interestingly, CXCR1 and CXCR2, the canonical receptors for CXCL1 and CXCL8, were not expressed in any cells in the atlas, whereas ACKR1, an alternative receptor for these ligands[62] was expressed solely on the vascular endothelial cells (Figure 5f). Cluster 16 showed increased signaling strength to the vascular endothelial cells through CXCL2-ACKR1 interaction (Fig 5e).

#### Changes in Cell Type Proportions with Sex

The atlas’ large sample size allowed us to evaluate possible associations between TME heterogeneity and sex, as the level and type of immune cell infiltration in a patient’s TME can affect their response to ICIs. Differential cell type compositional analyses were performed within the immune and non-immune compartments, separately.[63] Similar clusters were merged for simplicity, namely the naïve, Treg, CD4 Tcm and Tfh clusters were merged into one CD4 T cell cluster, the cytotoxic and dysfunctional CD8 T cells were merged into one CD8 T cell cluster, and the four salivary clusters were combined into one (Supplementary Figure 7a,b). Multiple significant changes were observed in the immune compartment, including an increased proportion in men of macrophages, CD8 and proliferating T cells, a cluster of myeloid cells (Mac/Mono/Neutrophils) and cDCs, and an increased proportion in women of B cells and Plasma cells (Fig 6a). A pan-cancer study looking at changes in cell type proportion using deconvoluted TCGA data also found CD8T cells to be increased in men, but a large number of other immune populations enriched in women.[64] In the non-immune compartment, the relative proportion of Epi1 and Epi4 increased in men, whereas Epi3 increased in women (Fig 6b). Equivalent analyses performed within the individual studies were largely unsuccessful in identifying significant sex-associated changes (Supplementary Figure 7c,d), highlighting once again the utility of the atlas in interrogating changes that require a large number of patients to achieve the necessary statistical power.

## Discussion

In this study, we integrated samples from six publicly available HNSCC scRNA-seq datasets to create the largest HPV-negative HNSCC atlas to date, providing a comprehensive characterization of the disease through both the contextualization of existing findings and the discovery of novel cell populations and their associations. The data are thoroughly annotated with patient metadata, cell types, and their signatures. The large number of patients enabled statistically powered evaluation of associations among gene expression, cell type composition, and molecular and clinical phenotypes.

As single-cell data are prone to batch effects that occur when experiments are performed in different labs, at different times, or using different instruments, integration methods that remove technical variability are necessary for clustering by cell type instead of by dataset.[65,66] Seurat’s reciprocal PCA method was found to effectively remove these confounding effects. Importantly, the rPCA method provides a corrected gene expression matrix that can be used for downstream clustering. After integration, we observed mixing of datasets and clear separation of cell types by canonical marker genes. Furthermore, cells annotated pre-integration as the same cell type were found to cluster together after integration.

Our analysis of the integrated immune compartment identified multiple subtypes of T cells including naïve CD4, Tfh, Tregs, CD8, and NK cells, which inhabited multiple states including cytotoxic, dysfunctional, and proliferating. Our annotation of the T cell compartment based on cytotoxic and dysfunctional state scores showed that T cells states fall on a trajectory, and that a single cell may simultaneously acquire characteristics of multiple states.[24,67] NK cells are generally considered cytotoxic and are associated with better prognosis in HNSCC.[68,69] CD8 T cells also exhibit this cytotoxic functionality in killing tumor cells. However, after prolonged stimulation by tumor antigens, CD8 T cells can become dysfunctional, also known as “exhausted”, a state associated with loss of cytokine production and effector killer function that is characterized by expression of immune checkpoint markers such as LAG3, TIGIT, CTLA4, and PD-L1.[70] Through the creation of an effector capacity score that considers both the levels of cytotoxicity and dysfunction of a single T cell, we show that while NK cells exhibit strong cytotoxicity, the CD8-positive T cells can be simultaneously enriched for both cytotoxic and dysfunctional scores, reflecting a multi-state phenotype that can be captured by the effector capacity score. Indeed, T cell cluster 0 exhibits this behavior, as it is enriched for both cytotoxic and dysfunctional signatures resulting in a lower effector capacity score, suggesting it may have started out as cytotoxic, but may have later acquired a more dysfunctional phenotype.

Additionally, we investigated the T cell effector capacity program’s association with tumor stage, as previous studies have shown changes in cell states can affect tumor progression across cancer types.[71–73] In HNSCC, higher numbers of CD8 T cells have been previously shown to be associated with better prognosis, but the state of the T cell was not considered.[74] We showed that T cells are most cytotoxic at stage I, and shift to a more dysfunctional phenotype at stage II as the tumors progress, possibly reflecting the ability of tumors to create immune suppressive microenvironments.

We then focused on the less characterized macrophage compartment, where we identified six subpopulations of macrophages and myeloid derived suppressor cells (MDSC). Two of the identified macrophage populations, SPP1 and CXCL9, recapitulate recent results that suggest the recasting of macrophage polarity along the SPP1 and CXCL9 axes rather than the traditional M1/M2 phenotypes, and associates them with worse and better survival, respectively.[34,75] We also identified a population of IL1β-positive myeloid cells not previously described in HNSCC that show both an immunosuppressive phenotype through the expression of IL1β and IL10, and an angiogenic phenotype, but lacking classical markers for M2 macrophages, and is SPP1-negative. Full characterization of this population may be beneficial for therapeutic intervention, as M2 macrophages are widely associated with decreased survival and are targeted by some therapies,[40] with a recent study also showing macrophage-secreted IL1β to promote drug resistance to docetaxel.[37] Through the integration, we were also able to identify a population of neutrophils which was not identified in any of the individual studies most likely due to low numbers, highlighting the ability of the atlas’ increased resolution to recover less-prevalent cell types.

The pooled analysis of epithelial tumor cells across datasets highlighted shared characteristics of these tumors and unique patient-specific niches. We identified a trajectory of differentiating epithelial cells. It’s been suggested that many cancers, including HNSCCs, maintain a population of cancer stem cells (CSCs) or tumor cells with stem-like properties, which can self-renew and give rise to differentiated cells.[76–78] Other plasticity-related signatures, such as epithelial-to-mesenchymal transition (EMT) and partial-EMT, have been described in which epithelial cells become more plastic and acquire mesenchymal features. We showed less differentiated cells (Epi1) to be enriched for stem-like properties, the pEMT signature, higher potency, and more likely to localize to the tumor edge and interact with stromal populations, recapitulating one of the major findings from Puram et al.[15] In contrast, differentiated cells (Epi4, Epi5) are more likely to inhabit the tumor core and less associated with stromal populations. We also identified patient specific epithelial subclones (Epi2, Epi3) enriched for xenobiotic metabolism and translation, respectively. Epi2 was also enriched for the NRF2 pathway which is activated by oxidative stress (cigarette smoke)[79], suggesting expansion of a specific subclone in an individual due to environmental effects, and importantly, highlighting that we can retain patient-specific populations through single-cell integration.

CAFs are a heterogeneous group of fibroblasts that promote tumor growth in several ways: through the remodeling of the extracellular matrix (ECM) to promote metastasis, through interactions with cancer cells to promote proliferation, and through the release of cytokines and chemokines that promote tumor invasion through the TME.[59,61,80] We identified seven clusters of CAFs, which we categorized into ECM-related CAFs, myofibroblasts, CXCL8 CAFs, and CXCL12 CAFs.

Because CAFs are in the early stages of being defined, it can be unclear how CAF populations identified in different studies are related.[81,82] For example, imCAFs are frequently referenced in the literature, with marker genes CXCL12, CXCL14, IL6, IL1, and IL8 (CXCL8)[80,81,83]. Through the atlas we identified two distinct imCAF populations, one defined by CXCL1/8, and another by CXCL12/14. This CXCL8 population also shares marker genes with the ‘resting’ fibroblast population initially described by Puram et al.[16]. The CXCL8 population has previously been identified and largely explored by Quah et al[27], who showed that it was induced by galectin-7 (LGALS7) expression in malignant epithelial clusters. We identify a novel potential interaction between CXCL8 CAFs and vascular endothelial cells through both cell-cell communication analysis and cluster co-occurrence analysis. A recent study which performed spatial transcriptomics on HNSCC showed coexpression of CXCL1, CXCL8 and ACKR1 in some patients.[51] Previous studies showed that CXCL8 and CXCL1 promote the angiogenic potential of endothelial cells and affect their proliferation and survival in-vitro.[84] A similar mCAF population also expressing CXCL8 and IL24 was recent identified in ESCC and found to facilitate metastasis and promote angiogenesis.[85] In HNSCC, both CXCL1 and CXCL8 have both been associated with poor patient survival.[76] Taken together, these results suggest that the CXCL8 CAF population may lead to worse outcome for HNSCC patients through stimulation of angiogenesis by vascular endothelial cells.

As sex-associated differences in cell type proportions have been explored in the context of some cancers but not in HNSCC, we used the atlas to address this question. The observed cell type proportion changes suggest men may be better candidates for ICI, as they have higher proportions of PDL1-expressing CD8 T cells in their TMEs, which has been shown to improve response to ICIs[86], corresponding to elevated proportions of proliferating and CD8T cells that are seen in men in the atlas (Fig 6a). Within the non-immune compartment, the Epi1 cluster, which is enriched for both the pEMT signature and proliferating tumor cells, was found to be enriched in men, consistent with a study that showed an increased rate of carcinogen-induced proliferation in male mice compared to female mice, leading to more aggressive and metastatic cSCC.[87]

As more HNSCC single-cells datasets become available, incorporating them into the atlas will improve characterization of cellular diversity at an ever higher resolution, and will provide increased statistical power to enable identification of additional associations. We offer the atlas as an integrated and thoroughly annotated resource for the characterization of HPV-negative HNSCC, hypothesis generation, and validation.

## Methods

### Data acquisition

Six datasets were integrated in this study, totalling 54 patients and 232,015 cells (Table 1). For five datasets, the processed matrix files containing raw counts and gene and barcode tsv files were downloaded from GEO: Peng[26] (GSE172577), Choi[23] (GSE181919), Cillo[24] (GSE139324), Quah[27] (GSE225331), and Kurten[25] (GSE164690). Due to availability, processed normalized data were downloaded for the last dataset, Puram[15] (GSE103322). HPV-positive samples were removed to retain only HPV-negative samples for use in the atlas.

### Dataset Pre-processing

Before integration, each dataset was pre-processed individually using *Seurat*[66] (RRID:SCR_007322) to eliminate low-quality cells and annotate the cells by cell type. If not pre-filtered in the originating study, cells with fewer than a minimum number of genes detected per cell (nFeatures ≤ 200) or more than a maximum percent of reads coming from mitochondrial genes (≥ 20%) were filtered out, as summarized in Table 3. These cutoffs were largely consistent with those chosen by the individual studies’ authors. We retained all cells for Puram, Quah, and Choi, which were already prefiltered.

**Table 2:** Processing Steps for pre-intervention.

| Dataset | Puram | Kurten | Cillo | Peng | Quah | Choi |
| --- | --- | --- | --- | --- | --- | --- |
| **Individual Dataset Pre-processing** |  |  |  |  |  |  |
| Step 1: QC |  |  |  |  |  |  |
| Filtering | Published data prefiltered | Filtered | Filtered | Filtered | Published data prefiltered | Published data prefiltered |
| nGenes/Cell | 2,000, or average housekeeping expression level < 2.5 | 200 | 200 | 200 | 200 | 200 |
| Percent mito | NA | 20% | 20% | 20% | 20% | 10% |

To annotate cells, each dataset was normalized by library size and log transformed using Seurat’s *NormalizeData* function, except for Puram, which was already normalized. Data was clustered using the Louvain algorithm with 2000 variable features and 30 PCs at a resolution of 0.3. Cells were annotated at a single-cell level using *SingleR*[88] (RRID:SCR_023120) with both the Blueprint[89,90,91] (RRID:SCR_003844) and Human Primary Cell Atlas (HPCAD)[91,92] databases, accessed through the *celldex* package.[91] Each cluster was assigned a broad cell label (Epithelial, Endothelial, Fibroblast, T/NK, Immune, B Cell, or Mast), based on the consensus majority label amongst Blueprint and HPCAD, and when available, the paper-provided annotations.

### Integration

Integration was performed via Seurat’s reciprocal principal component analysis (rPCA) method.[93] For each pre-processed dataset, *SCTransform* was used to normalize the raw counts using *method = “glmGamPoi”*, except for Puram, in which the already normalized counts were used. *SelectIntegrationFeatures* was used to identify 3,000 integration features (genes highly variable amongst all datasets). The *FindIntegrationAnchors* function was used to identify 20 anchors in 30 dimensions. The integration was then performed using *IntegrateData*, producing a single Seurat object containing two slots; an integrated slot which contains integrated-corrected counts for the 3000 integration features, used only for clustering, and an RNA slot containing the raw counts for all genes. The raw data in the RNA slot was normalized by library size and log transformed using *NormalizeData*, as well as saved to a new SCT slot in which it was normalized using *SCTransform*, with *vars.to.regress = ‘dataset’* (again, the pre-normalized Puram data was copied into the SCT slot and not renormalized (Supplementary Fig 1)). After integration, dimensionality reduction and clustering were performed on the integrated slot using 30 PCA dimensions, using the Louvain algorithm, at a resolution of 0.4.

### Signature scoring

Transcriptional signatures were collected from multiple sources (Table 4). Szabo[30] and Chu[94]: pan-cancer derived T cell signatures. Luen[67]: a review finding commonalities between T cell signatures across papers. Mulder[95]: monocyte and macrophage cross-tissue derived signatures. Puram[16]: epithelial cell states derived from HPV-positive and HPV-negative scRNAseq, and cell type signatures. Cheng[96]: pan-cancer tumor infiltrating myeloid cell derived signatures. Cho[59]: pan-cancer derived fibroblast signatures. Select genesets from the Kegg[97] (RRID:SCR_012773), Hallmarks[98], and Reactome[99] (RRID:SCR_003485) compendia were also used. Signatures of HNSC tumor core versus leading edge were compiled from Arora et al.[51]

**Table 4:** Signatures used in atlas.

| Publication | Derivation | SignatureType |
| --- | --- | --- |
| *Szabo et al.* | Pan-cancer | T cell |
| *Chu et al.* | Pan-cancer | T cell |
| *Luen et al.* | Pan-cancer | T cell |
| *Mulder et al.* | Cross-tissue | Myeloid |
| *Puram et al.* | HPV+, HPV- HNSCCs | Epithelial states and TME cell types |
| *Cheng et al.* | Pan-cancer | Myeloid |
| *Cho et al.* | Pan-cancer | Fibroblast |
| *Kegg* |  | General |
| *Hallmarks* |  | Tumor |
| *Reactome* |  | General |

For all gene expression analyses, the normalized data slot of the RNA assay was used. In each compartment, cells were scored for a given signature using Seurat’s *AddModuleScore*.[100]

### Creation of effector cell score and correlation with stage

All T cells were scored using the dysfunctional and cytotoxic signatures from Luen et al[67]. An effector capacity score was created by subtracting the dysfunctional score from the cytotoxic score for each cell, so that the higher the score, the more purely cytotoxic the cell is assumed to be. The relationship between the effector capacity score and stage was determined using a mixed effect model using *lme4*[101] (RRID:SCR_015654), with dataset and patient as nested random effects.

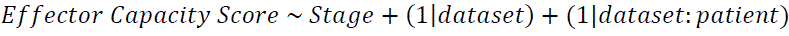

The association of effector capacity score with stage was also performed separately within each dataset using the same model, but with just patient as a random effect.

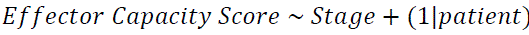

### Identification of marker genes

Identification of marker genes per cluster was performed using Seurat’s *FindAllMarkers* with the parameters *test.use* = ‘MAST’ (RRID:SCR_016340), and *latent.vars* = ‘dataset’ to control for dataset as a fixed effect[102].

### Gene set enrichment analyses

Gene set enrichment analyses were performed by hypergeometric testing with the Kegg[97] (RRID:SCR_012773), Hallmarks[98], Wikipathway[103] (RRID:SCR_002134), and Biocarta[104] (RRID:SCR_006917) compendia using hypeR[105], with all genes in the atlas used as the background.

### Taxonomic organization of cell type clusters

K2Taxonomer[33] was used to elucidate the relationship between different clusters and define transcriptional programs enriched at data-driven cluster subgroups. K2Taxonomer was applied separately to the immune and the epithelial compartments, using the scaled, integrated slot of the dataset. The preprocessing step was performed using *featMetric=“F”*, *nBoots=400*, and *clustFunc=cKmeansWrapperSubsample*. The K2 dendrogram was produced using the main *K2tax* function with default settings. The normalized RNA slot data was used to perform downstream differential expression and gene set enrichment analyses, with the dataset source included as a covariate.

### Cluster co-occurrence analyses

Cluster co-occurrence analyses were performed on the immune and non-immune compartments separately to identify clusters manifesting coordinated proportional changes across patients. Per patient cluster proportions were estimated as the number of cells in a cluster divided by the total number of cells in the given compartment. Spearman correlation was calculated among all clusters across all patients. Correlations with a p-value < 0.05 and coefficient greater than 0.01 were considered significant.

### Inference of copy number variations

InferCNV[106] (RRID:SCR_021140) was used to detect copy number variations (CNVs) in the epithelial cells, and was run on each dataset separately. Epithelial cells were considered cancer cells and used as ‘observations’. A random set of 1,000 Fibroblasts and T cells was sampled and used as reference cells. As a negative control, a subset of 100 of the 1,000 reference cells was removed and added to the observation set of epithelial cells in which CNVs should not be found. The gene position file hg38_gencode_v27.txt was used for gene locations. Raw counts were used as input, and inferCNV was run using *cutoff = 0.1*, *cluster_by_groups = TRUE*, *denoise = TRUE* and *HMM = FALSE*.

The output of InferCNV is a gene-by-cell matrix containing CNV scores centered at one, where a value less than one corresponds to a loss in copy number and a number above one corresponds to a copy number gain. A malignancy score for each cell was calculated by subtracting one from the copy number score to center at zero, taking the absolute value of the sum of scores per cell, and dividing by the number of genes.

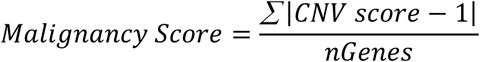

### Inference of cell potency

CytoTrace2[48] was run separately on each dataset, and on the immune and non-immune compartments separately. The counts slot of the RNA assay was used as input. The final potency annotation was added to the atlas for visualization across datasets.

### Detection of cell type proportion changes across sex

Sccomp[63] was used to detect changes in cell type proportions between males and females. Sccomp was applied to the immune and nonimmune compartments of the atlas separately to avoid artefacts arising from CD45+ and CD45-cell being sorted and sequenced separately which was the case in many of the individual datasets. Sccomp was first applied to each compartment using all datasets in the atlas, and controlling for dataset as a random effect in both the composition and variability models with bimodal_mean_variability_association = TRUE. Sccomp was then applied to the immune and nonimmune compartments on each dataset separately using the same parameters, but without including dataset as a random effect.

### Cell cell communication analyses

CellChat[107] (RRID:SCR_021946) was used to perform cell cell communication analyses on the RNA slot between clusters as specified in Supplementary Figures 7 a and b. The communication probability pathway was computed using the “trimean” method, requiring a minimum of 10 cells. A second CellChat analysis focused on signaling between the specific myeloid clusters identified in Figure 3 was performed which used the clustering specified in Supplementary Figure 7a and b, but in which the macrophage and mac/mono/neutrophil clusters were resolved at a higher resolution as detailed as the six clusters in Figure 3. Lastly, a CellChat analysis focused on signaling occurring between the fibroblast populations identified in Figure 5 was performed using the clustering shown in Supplementary Figure 7 and b, but in which clusters Fib1,Fib2, and Fib3 were resolved at the higher resolution of 0.7 as shown in Figure 5a.

## Statistics

Confidence intervals were used for all linear regression models. An fdr-corrected hurdle p-value was used for all differential expression testing through MAST (RRID:SCR_016340). For cluster co-occurrence analyses, spearman correlations were used and correlations with a p-value < 0.05 and coefficient greater than 0.01 were considered significant.

## Availability of Data, Source Code and Materials

The full atlas will be available as a Seurat .rds object available for download on zenodo upon publication, and has been deposited to cellxgene for online visualization, also public upon publication.

## Declarations / Acknowledgements and Competing Interests

The authors declare no competing interests.

## Consent for publication

Not applicable. All datasets included are already published.

## Funding

This study has received funding from:

System-Level Analyses of Multi-Omics Data to Reveal Mechanisms of Head & Neck Cancer; R01DE031831; National Institute of Dental and Craniofacial Research

Defining immune-evasive mechanical signaling in head and neck cancer.; R01DE033519; National Institute of Dental and Craniofacial Research

Elucidating mechanisms of cellular communication critical for head and neck cancer progression and metastasis.; 1F31DE033292-01; National Institute of Dental and Craniofacial Research

NIH/NCATS grant BU-CTSI 1UL1TR001430 (SM)

## Author contributions

L.K. performed all analyses and wrote the manuscript. A.C. and E.R. provided computational guidance and feedback. A.S, M.K. and X.V. provided biological guidance. S.M. oversaw the project, provided feedback and guides the research. The manuscript is edited by all authors.

## List of Abbreviations

HNSCC: Head and Neck Squamous Cell Carcinomas
HPV: Human papilloma virus infection
EMT: Epithelial-to-mesenchymal transition
pEMT: partial EMT
CAFs: cancer-associated fibroblasts
NK: Natural killer
MDSCs: myeloid-derived suppressor cells
scRNA-seq: Single-cell RNA-sequencing
HPCAD: Human Primary Cell Atlas Database
rPCA: reciprocal principal component analysis
CNVs: copy number variations
Tfh: T follicular helper cells
Tregs: regulatory T cells
Tcm: central memory T cells
CSCs: cancer stem cells
FDR: False discovery rate
ECM: Extracellular Matrix

---

**Supplementary Figure 1:**
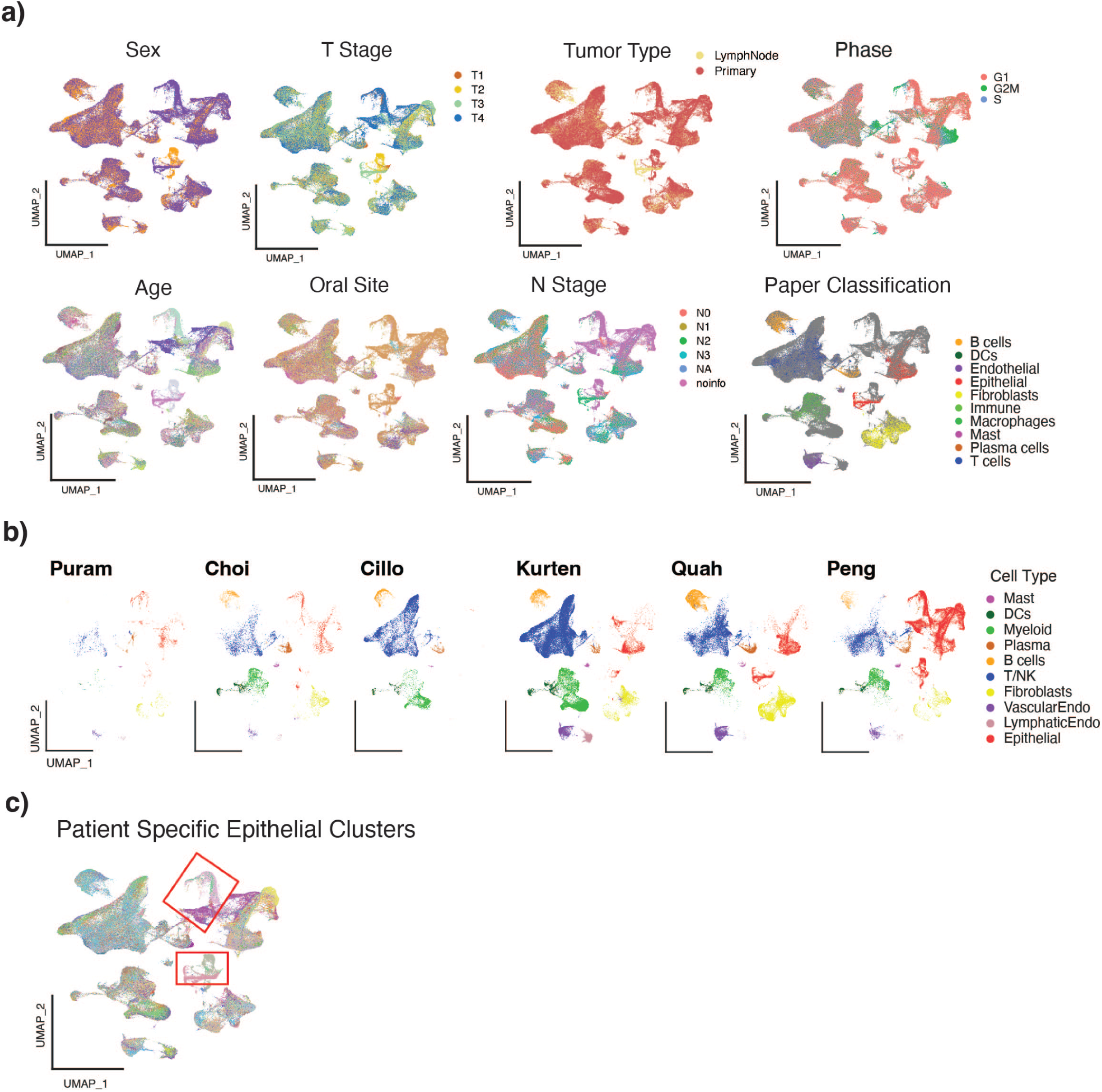
Summary of Atlas Metadata. a) Visualization of metadata on atlas UMAP. b) Integrated UMAP split by dataset and colored by cell type. c) Atlas UMAP colored by patient. Red boxes highlight epithelial or salivary clusters with most cells originating from one patient.

**Supplementary Figure 2:**
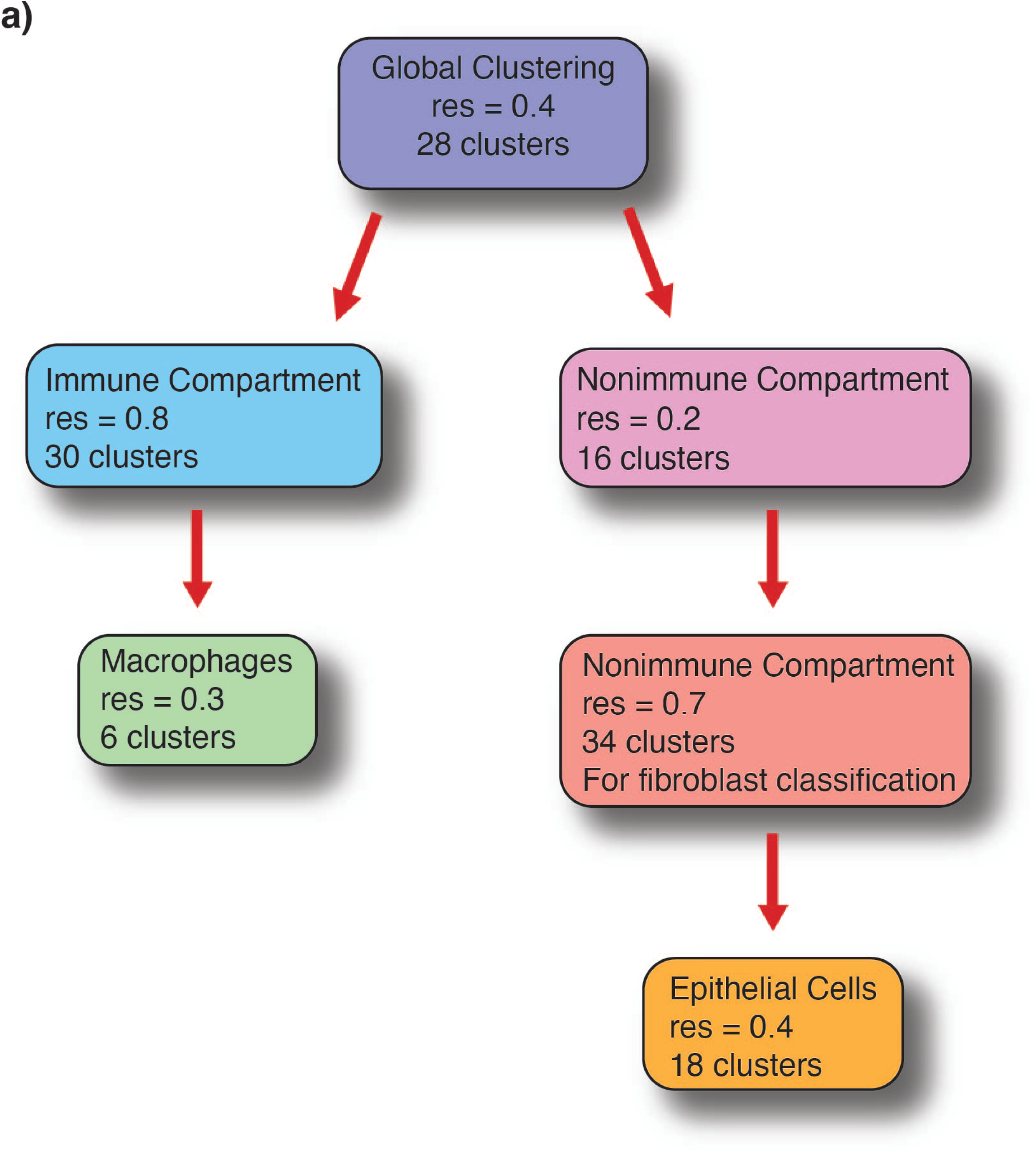
Overview of clustering performed in different compartments at various resolutions and number of clusters found.

**Supplementary Figure 3:**
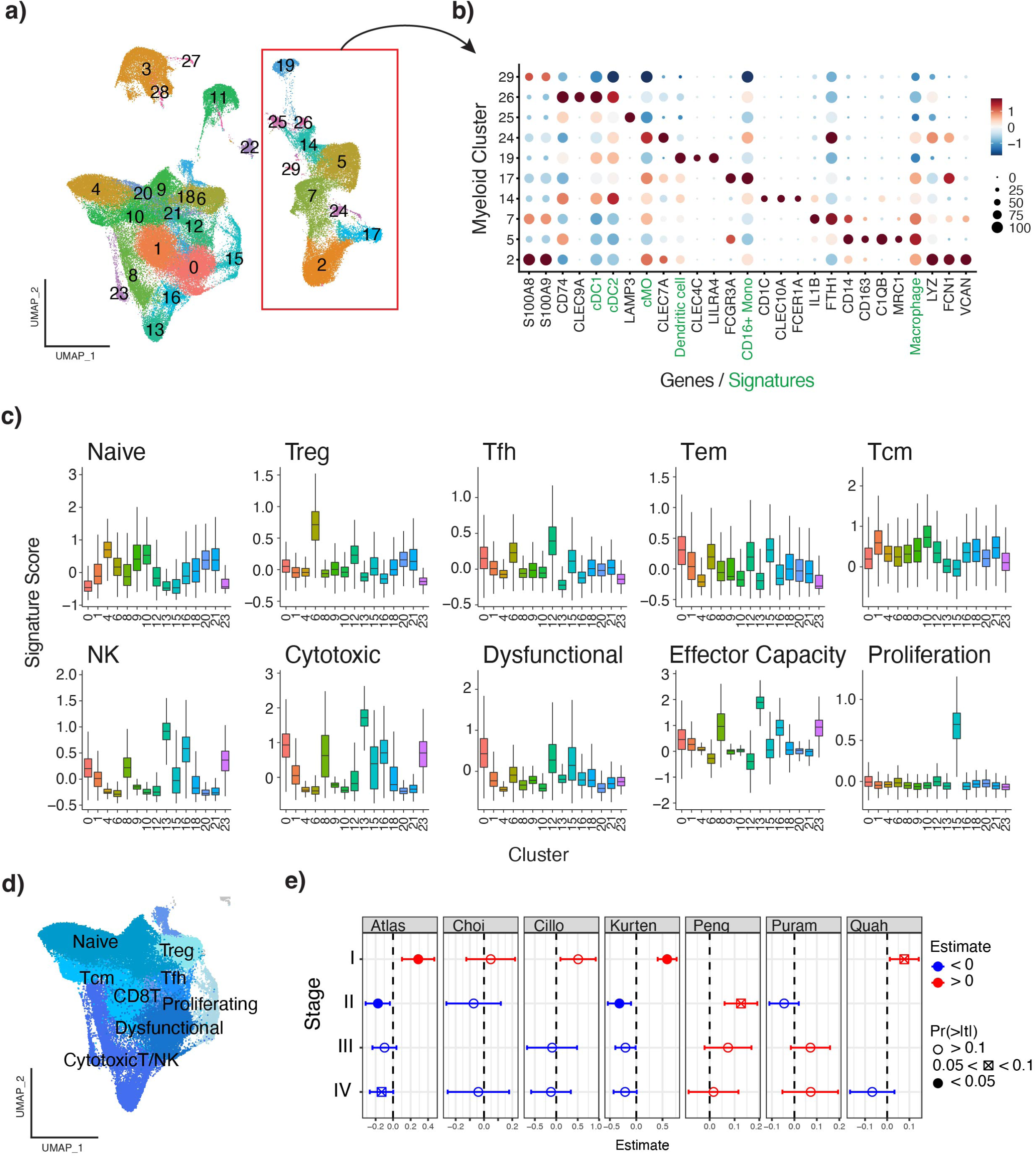
Identification of T cell and myeloid cell subpopulations and association with stage. a) UMAP of the immune compartment clustered at resolution 0.8. b) Dotplot of marker genes (black) and signatures (green) enriched in the myeloid clusters. c) Boxplots showing the module score of different T and NK cell signatures across the T and NK cell clusters. d) Summary of T cell classification based on signature enrichment of the T cells. e) Estimates for the association of T cell effector capacity score with stage across all datasets in the atlas (left), followed by within each dataset separately.

**Supplementary Figure 4:**
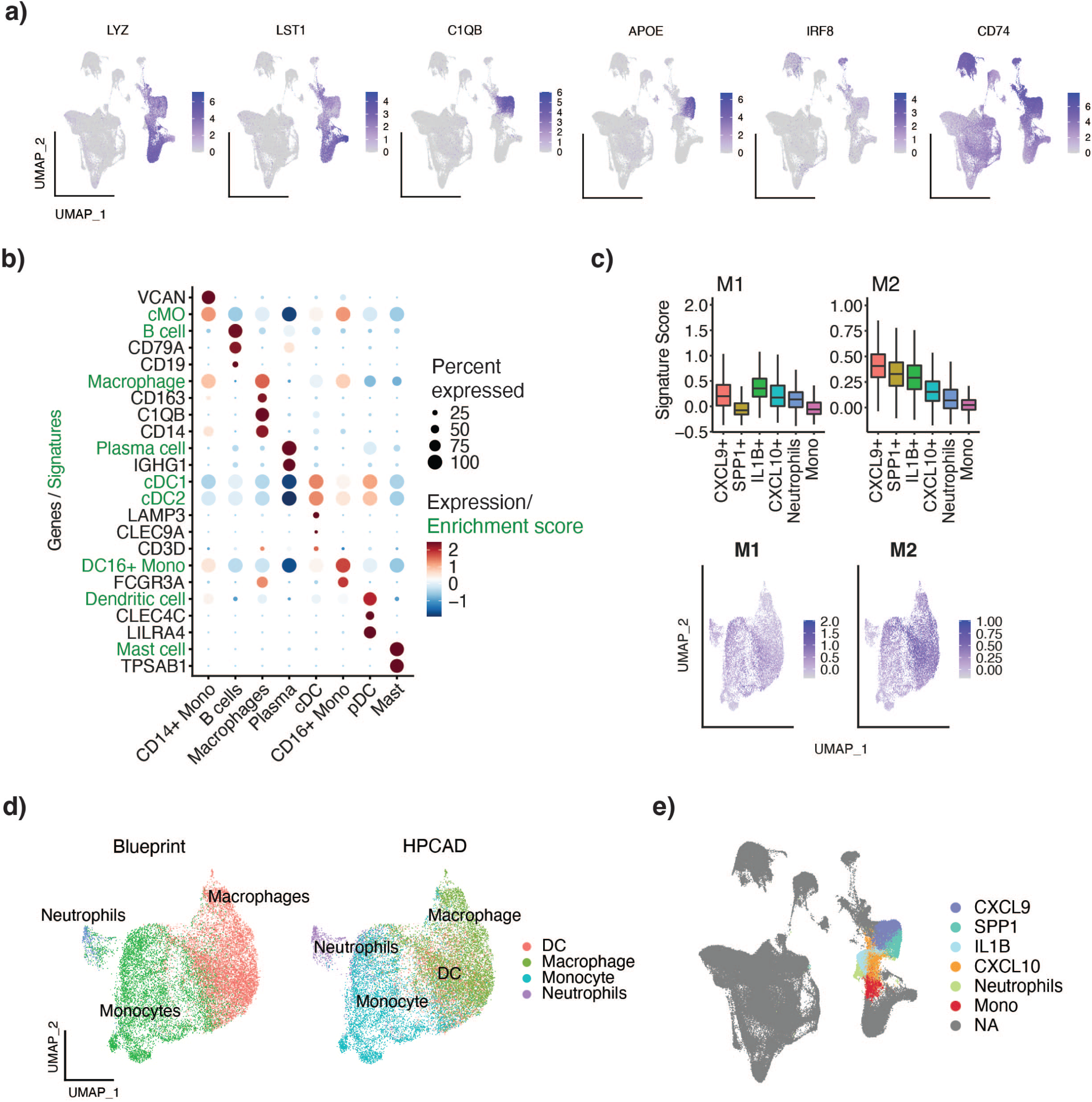
Macrophage cell classification. a) UMAP of the immune compartment showing genes influential in myeloid K2 Taxonomer clustering. From left to right: LYZ, LST1, C1QB, APOE, IRF8, CD74. b) Signature (green) and gene (black) expression in myeloid clusters. c) M1 and M2 signature score in macrophage subclusters. d) Blueprint and HPCAD cell classifications of top cell types identified. Supplementary table 3 contains all classifications. e) Immune compartment UMAP colored by myeloid sub-clusters identified in Fig 3.

**Supplementary Figure 5.**
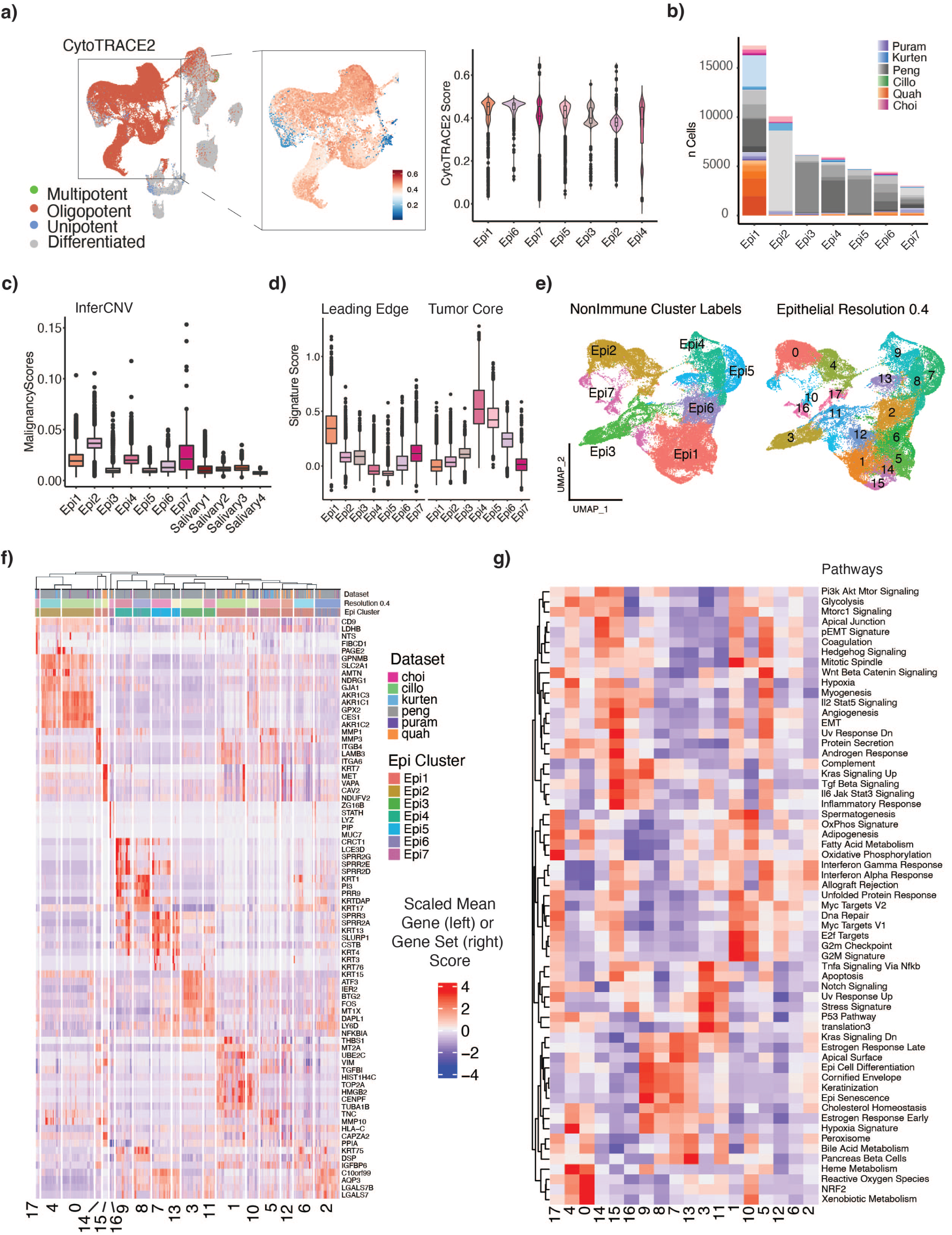
High-resolution characterization of the epithelial cells. a) CytoTRACE2 potency classification (left) across epithelial clusters, and scores (UMAP middle and violin plots right) across epithelial clusters. b) Number of cells per patient per cluster. Each dataset is a color, which each patient a different shade within the color group. c) InferCNV malignancy scores across clusters. d) Boxplots showing enrichment scores of Arora et al.’s leading edge and tumor core signatures across epithelial clusters. e) Epithelial cells reclustered and labelled by initial whole-immune compartment clustering (left) and by higher resolution 0.4 clustering (right). f) Heatmap of top five genes per cluster at resolution 0.4. g) Heatmap of enrichment scores of Hallmark genesets and signatures from Puram et al. (2017) per each cluster found at resolution 0.4.

**Supplementary Figure 6.**
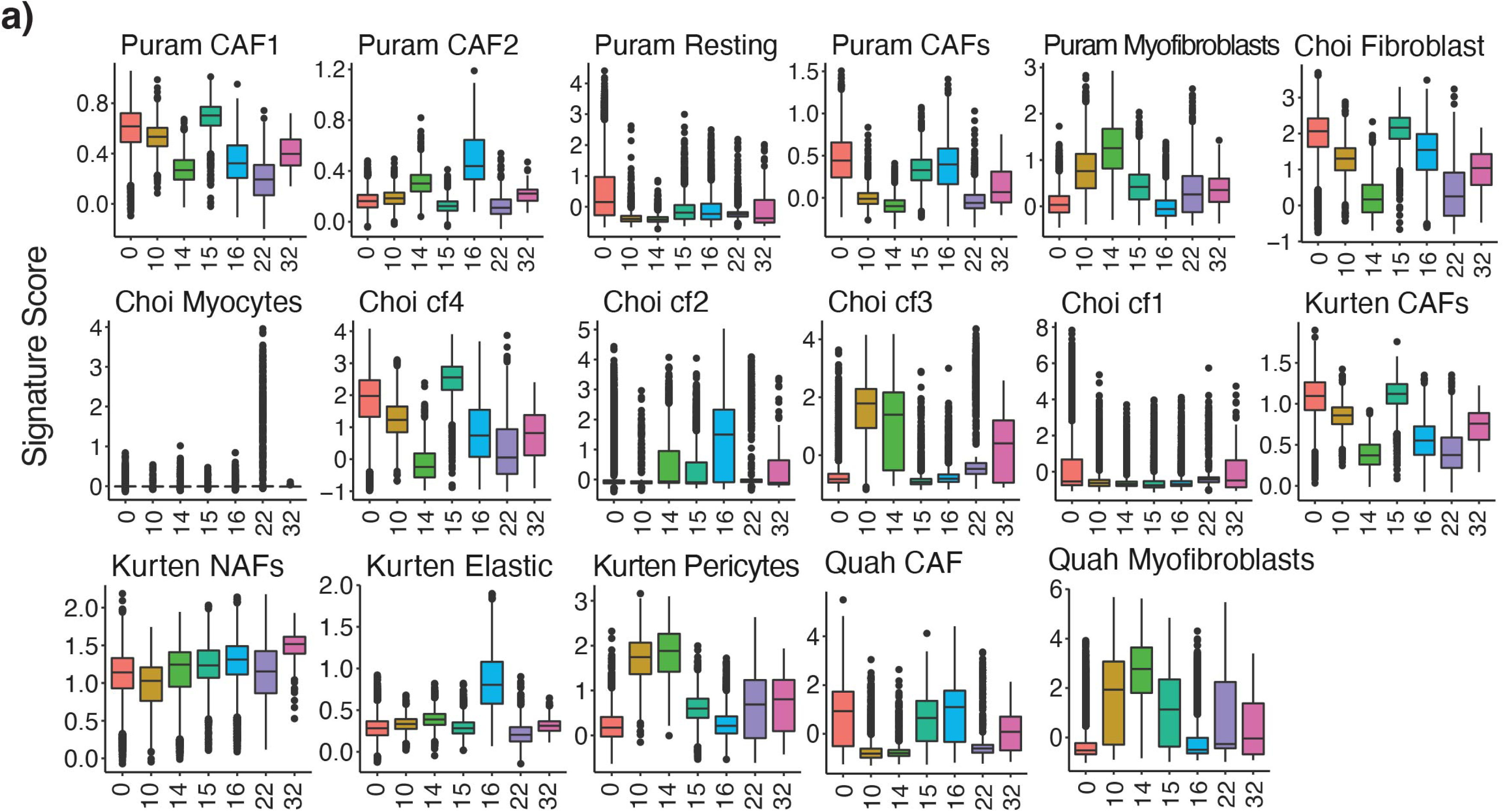
Reconciliation of fibroblast terminology. a) Enrichment of fibroblast signatures from original publications in seven fibroblast clusters.

**Supplementary Figure 7.**
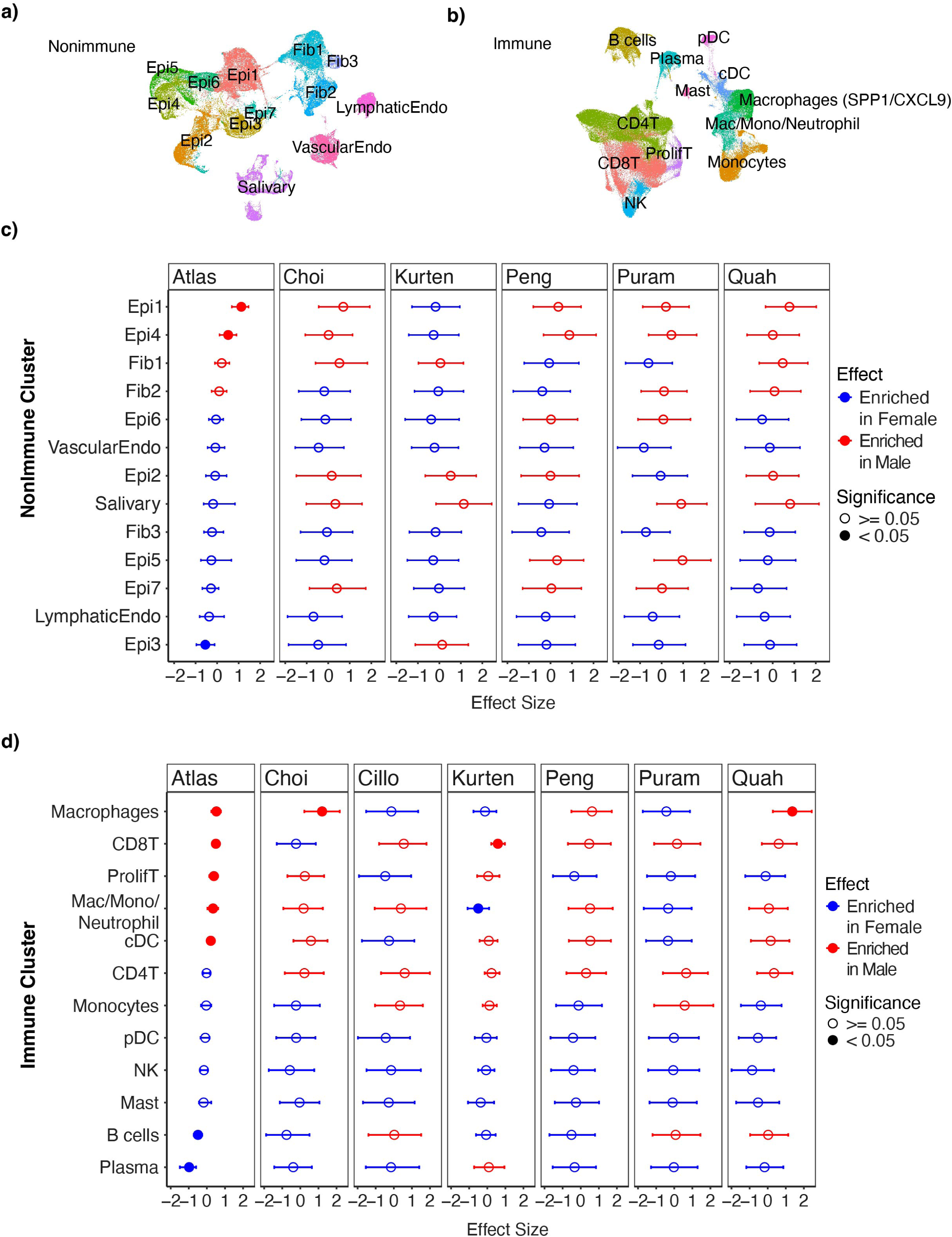
Sex-specific cell type proportion changes on a dataset level. Clustering in the a) nonimmune and b) immune compartments used as input to sccomp. c) Cell type proportion changes identified by sccomp within the nonimmune compartment when using the whole atlas (left), and individually within each dataset. d) Cell type proportion changes identified by sccomp within the immune compartment when using the whole atlas (left), and individually within each dataset.

